# Mating preferences act independently on different elements of visual signals in *Heliconius* butterflies

**DOI:** 10.1101/2023.11.24.568567

**Authors:** S. H. Smith, L. Queste, D.S. Wright, C.N. Bacquet, R.M. Merrill

**Affiliations:** Division of Evolutionary Biology, LMU Munich, Grosshaderner Str. 2, 82152 Planegg-Martinsried, Germany; Universidad Regional Amazónica de Ikiam, Tena, Ecuador; Smithsonian Tropical Research Institute, Gamboa, 0843-03092, Panama

## Abstract

Mating cues are often comprised of several elements, which can act independently, or in concert to attract a suitable partner. Individual elements may also function in other contexts, such as anti-predator defense or camouflage. In *Heliconius* butterflies, wing patterns comprise several individual color pattern elements, which advertise the butterflies’ toxicity to predators. These wing patterns are also mating cues, and males predominantly court females that possess the same wing pattern as their own. However, it is not known whether male preference is based on the full wing pattern or only individual pattern elements. We compared preferences of male *H. erato lativitta* between female models with the full wing pattern and those with some pattern elements removed. We found no differences in preference between the full wing pattern model and a model with pattern elements removed, indicating that the complete composition of all elements is not essential to the mating signal. Wing pattern preferences also contribute to pre-mating isolation between two other *Heliconius* taxa, *H. erato cyrbia* and *H. himera*, therefore, we next compared preferences for the same models in these species. *H. erato cyrbia* and *H. himera* strongly differed in preferences for the models, potentially providing a mechanism for how pre-mating isolation acts between these species. These findings suggest that contrasting levels of selective constraint act on elements across the wing pattern.

## Introduction

Animal signals are highly diverse and often involve multiple elements, acting across various sensory modalities. Even within a single modality, different elements may target different receivers and/or communicate different information (Candolin 2003). For example, visual cues can include multiple discrete color and pattern elements (Endler 1978), as well as ornamentation or movement (Fleishman 1988). These signals may be shaped by several sources of selection acting in conflict or co-operation, including predation and sexual selection. Determining how receiver behavior responds to individual signal elements can therefore help us understand how the evolution of composite signals is constrained and/or contributes to diversification.

Incorporating multiple elements can enhance signaling function in several ways. Certain elements may be more effective when viewed in unison (Guilford & Dawkins 1991, Brooks & Caithness 1995). For example, in budgerigars (*Melopsittacus undulatus*) the high contrast generated by adjacent UV fluorescent and reflective plumage patches is more attractive to mates than either element in isolation (Arnold et al. 2002, Hausmann et al. 2003). Composite signals may also allow transmission over a broader range of sensory environments. Male *Schizocosa* wolf spiders use both vibratory and visual courtship signals, which they adjust to more effectively signal across different sensory backgrounds (Hebets & Papaj 2005, Gordon & Uetz 2011). Finally, individual elements can carry independent information. In American goldfinches (*Carduelis tristis*), carotenoid-based plumage and bill coloration transmit information about parasite load, whereas the melanin-based black cap signals social status (McGraw & Hill 2000).

Despite this, it is increasingly apparent that mate preferences are sometimes based on only a subset of the total signal elements. Male *Eleutherrodactylus coqui* tree frogs, for example, produce a distinctive two-note call to communicate with conspecifics, but females respond to only the second note (Narins & Capranica 1976). Similarly, female preferences in mallard ducks primarily depend on bill color brightness, even though males display various other visual ornaments (Omland 1996). A study of male preference in Lycaenid butterflies showed that preference is based on an apparently very minor part of the overall signal. By subtly manipulating the wing patterns of female *Lycaenides melissa* to change the size of the hindwing spot and the associated ‘aurorae’ to mimic those of its sister species, *L. idas*, Fordyce et al. (2002) found that male *L. idas* preferred the manipulated wing patterns, even though the rest of the wing matched the heterospecific pattern. Together these examples highlight that different visual elements may serve different evolutionary functions, and perhaps that selection may not be uniform across different elements.

Numerous studies have now shown that traits under divergent ecological selection are additionally used as mating cues (Servedio et al. 2011). These types of traits may be important during speciation, because they couple divergent selection acting on the ecological trait to mating preferences, thereby promoting the evolution of premating barriers (Gavrilets, 2004; Servedio 2011). Warning patterns, advertising toxicity or unpalatability to potential predators, may act in this way if they are additionally used as mating cues by conspecifics. For example, the bright warning signals of the toxic strawberry poison dart frog, *Oophaga pumilio*, are used also used by females during mate choice (Summers et al. 1999, Siddiqi et al. 2004).

A key feature of aposematic signals is visibility and recognizability, which is often achieved by patterns of adjacent color patches, contrasting strongly both with each other and the background scene (Stevens & Ruxton 2012). This may imply that the function of aposematic cues relies on them being perceived in composite, while elements of mating cues can act independently from each other (however, aposematic signal function can also be based on only a single pattern element, see Winters et al. 2017). Given the multi-component nature of aposematic signals, it is unclear how mate preference acts when the same elements that make up aposematic signals are also used as mating signals.

The Neotropical *Heliconius* butterflies exhibit a huge diversity of bright warning patterns, which are often associated with Müllerian mimicry (Merrill et al. 2015). These warning patterns are both under strong positive frequency dependent selection (Mallet & Barton 1989; Merrill et al 2012), and are also used as mating cues (Crane 1955, Jiggins et al. 2001). Specifically, multiple studies have now shown that *Heliconius* males normally prefer to court females that share their own color pattern over those of closely related taxa, (e.g. Jiggins et al. 2001, Jiggins et al. 2004, Estrada & Jiggins 2008; Merrill et al. 2014), and this contributes to a strong premating reproductive barrier between closely related species (Jiggins et al. 2001). The extremely close mimicry exhibited between the distantly related species *H. erato* and *H. melpomene* suggests that predators discriminate against deviations in pattern across the entire wing. However, although both *H. erato* and *H. melpomene* males are known to discriminate between females based on visual cues (Jiggins et al. 2004, Estrada & Jiggins 2008) relatively little is known about which pattern elements trigger the behavioral response. Previously, Estrada & Jiggins (2008) compared preferences between geographical populations of *H. erato* with divergent warning patterns and found that males were more likely to be attracted towards females with at least one of the wing pattern elements in common to their own, over females with more different warning patterns. Additionally, Jiggins and colleagues (2001) found little difference in the response of *H. melpomene* to visually near identical models made from females from populations with or without a yellow hindwing bar. Finkbeiner et al. (2014) also showed that color is a more important determinant of male attraction towards *H. erato petiverana* females than the arrangement of these colors (i.e. pattern). However, no experiments have explicitly tested whether the whole composite wing pattern elicits male courtship behaviors, or just specific elements.

To determine how visual preferences are influenced by individual elements of composite warning patterns, we manipulated individual elements of the *H. erato lativitta* color pattern and tested the corresponding male preferences. To understand how preference for pattern elements could impact divergence of populations, we also investigated how males from two other Erato clade populations, *H. himera* and *H. erato cyrbia*, responded to manipulated *H. erato lativitta* wing patterns. These two sister taxa inhabit different habitats and differ in wing pattern, but maintain a contact zone in which hybridization occurs at a rate of 10% (Jiggins et al., 1996). Experimental evidence has shown that isolation is mainly driven by visual preference differences; both prefer their own color pattern over each other’s and do not exhibit any post-mating reproductive isolation (Merrill et al., 2014). Therefore, they provide an interesting case to investigate whether intraspecific mating can be maintained by differences in individual elements of the wing pattern, particularly within a conspecific background. Overall, our results suggest that not all elements of the aposematic pattern are important for mate choice, and that differences in preference based on individual pattern elements could play a role in the divergence of closely related species.

## Methods

### Butterfly collection and rearing

All experiments took place in insectaries at Universidad Regional Amazónica Ikiam, Tena, Ecuador (Figure 1, red diamond) between November 2021 and January 2022. *H. erato lativitta* were collected in the southeast of Napo province, Ecuador, whereas *H. erato cyrbia* and *H. himera* came from insectary populations established from individuals collected in southwest Ecuador (Figure 1). Butterfly stocks were maintained in separate 2 x 2 x 2m cages for each sex and species, with 10-50 individuals per cage. Each cage was supplied with nectar-bearing flowers, and ∼20% sugar solution with supplementary pollen for additional nutrition (Gilbert 1972), and *Passiflora punctata* shoots for oviposition in female cages. Caterpillars from these stocks were reared indoors and fed *ad libitum* on *Passiflora punctata*.

**Figure 1.**
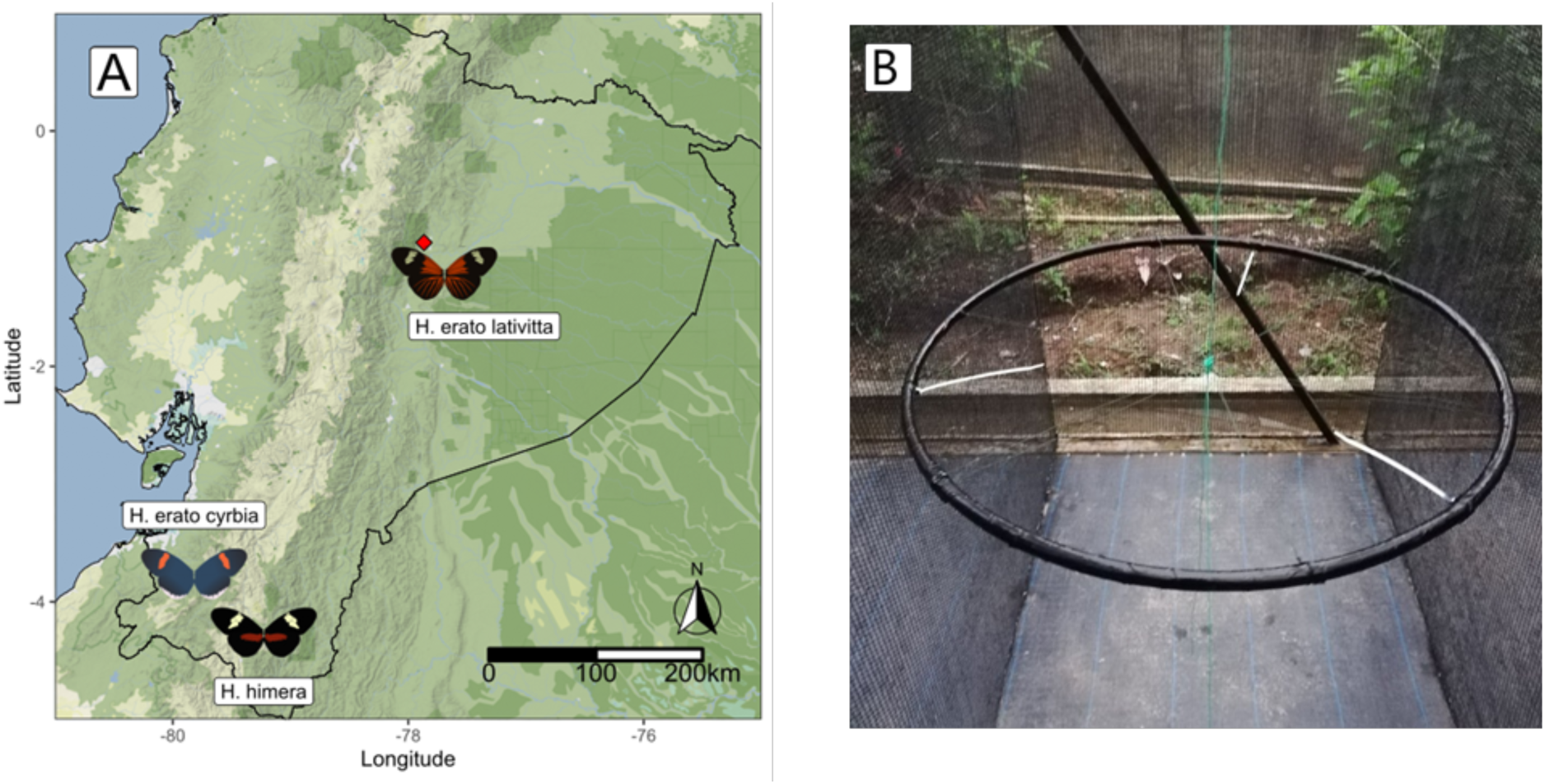
(A) Topographical map of Ecuador, showing the Andes mountain range in the center. Experiments took place at Universidad Regional Amazónica IKIAM (red diamond) and butterflies were collected at the sites surrounding the locations marked with images of the butterflies. (B) Layout of experimental cage, with circular frame suspended in the center. The three models with manipulated wing patterns of H. erato lativitta females were attached to the ends of the white sticks extending inwards from the frame. Preference behaviors of males of *H. erato lativitta, H. erato cyrbia* and *H. himera* were observed from outside the cage.

### Mounted female wing models

To make the female wing models, wings from sacrificed *H. erato lativitta* females were removed and subsequently washed for 5 minutes in dichloromethane to remove pheromones (Darragh et al. 2017). The wings were then manipulated by coloring over a patch of color on both sides of the wing with a black marker pen (COPIC Ciao Black 100 (Too Marker Products Inc.)) to generate one of three distinct wing patterns: ‘intact’, ‘red-only’ or ‘yellow-only’ as shown in Figure 2A-C. The remaining unmanipulated part of the wing was colored with a colorless marker pen with the same solvent (COPIC Ciao Colorless Blender 0 (Too Marker Products Inc.). Wings were then glued to white card, using a thin paper strip to allow flapping motion, mounted onto a black foam strip that acted as the butterfly’s body and allowed to dry for at least 2 hours before use.

**Figure 2.**
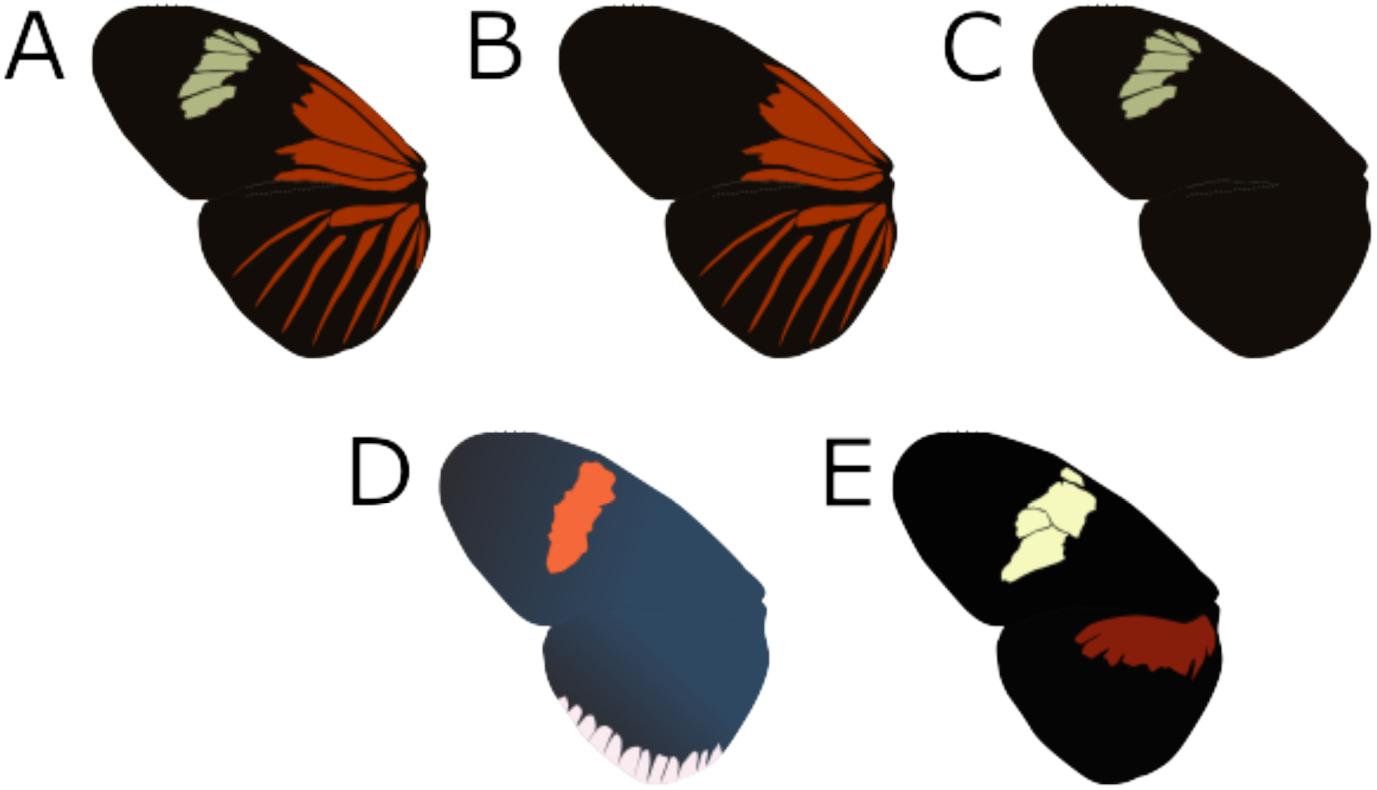
(A – C) Model types used to test mate preference based on wing pattern elements. (A) the full *H. erato lativitta* wing pattern, used for the ‘intact’ model; (B) ‘Red only’ model, featuring only the red dennis-ray elements of the *H. erato lativitta* wing pattern; (C)‘Yellow only’ model, featuring only the yellow forewing band of the *H. erato lativitta* wing pattern. (D) *H. erato cyrbia* wing pattern. (E) *H. himera* wing pattern. Figures are a graphical representation of the model types or wing patterns. Figure shows only the left dorsal wing pattern, but patterns were identical on all sides of the wing.

### Male preference experiments

Our experimental design involved assessing the preference of male *H. erato lativitta*, *H. erato cyrbia* and *H. himera* between models of *H. erato lativitta* females with three types of wing pattern manipulation. Trials took place between 08:00 and 17:00, and males were tested at least 5 days after eclosion. One male each of *H. erato lativitta*, *H. erato cyrbia* and *H. himera* were tested together in the same trial, and the combination of three males tested together was randomized between trials. No interactions between males that would impede a male’s ability to interact with the models were observed. Males were released into experimental cages at between 1 and 24 hours before the trial to acclimatize, during which they were provided with nectar-bearing flowers and sugar water solution, as above, but these were removed during the trial.

During 30-minute trials, males were presented with a set of three female models (intact, red-only and yellow-only) mounted on a circular frame, 90cm in diameter, suspended 1m from the ground (Figure 1B). The models were randomly placed at one of three positions on the frame, equally spaced at 60cm away from each other. A string attached to the center of the frame was pulled continuously throughout the trials to simulate wing movement (Klein & de Araújo 2010). Four different sets of models were used to account for individual variation between models. At the beginning of each trial, we recorded the date, time, weather, (recorded as ‘clear’, ‘under 50% cloud cover’, ‘over 50% cloud cover’ or ‘overcast’), model set and temperature.

Two types of behavioral interactions between the males and the models were recorded during the trial, using the Field Notes app (Neukadye). These were ‘approach’ and ‘courtship’. We defined ‘approach’ as a male directing its flight towards and coming within 10cm of the model. ‘Courtship’ was defined as ‘approach’ but additionally included hovering above the model for at least 1 second. Accordingly, if a male courted the model, this was also recorded as ‘approach’. Both behaviors were scored independently once per minute, for each model type and male, and counts of each behavior per trial were used in the subsequent analysis.

Preference trials for each male were repeated 5 times for each model set to account for individual variation and differences in male activity levels between different day times and weather conditions. Each male was tested on multiple model sets when possible, to account for variation in female model quality, though this was not possible for cases in which males died between tests on different sets, therefore most males (60%) were tested with only one set. Each model set was used for testing at least two different males of each species. Males were tested a maximum of 2 times per day and remained in experimental cages between the 5 repeated trials per model set, in the presence of other males.

### Statistical analysis

To determine whether males responded differently to the three model types, we fitted generalized linear mixed models (GLMM) with Poisson error distribution, implemented in the lme4 package (Bates et al. 2015) in R (R Core Team 2021). The dependent variable was either ‘approach’ or ‘courtship’ encoded as counts of minutes per trial in which a male did the behavior towards a model. Saturated models included male species, model type and their interaction as dependent variables. Weather was also included as a fixed factor with four levels: ‘100% overcast’, ‘>50% overcast’, ‘<50% overcast’ and ‘clear’ which correlated strongly with temperature (see supplementary materials). Male ID and trial nested within model set were included as random variables in all models to account for repeated measures. For some models tested, we implemented model optimizer BOBYQA (Powell 2009) to overcome convergence warnings associated with the complex structure of the models. When necessary, we subsequently checked for deviation between log likelihoods of different optimizers using the command allFit from the lme4 package (SM Table 1). Significance of fixed factors was determined using likelihood ratio tests (LRTs) by comparing models with and without individual terms. Effect sizes were calculated as the estimated marginal means (EMM) of the minimum adequate model using the emmeans package (Lenth 2021). We generated ternary plots of the raw data to visualize differences in ‘approach’ and ‘courtship’ by each male type, towards the three models. All analysis scripts are included in the supplementary materials.

**Table 1.**
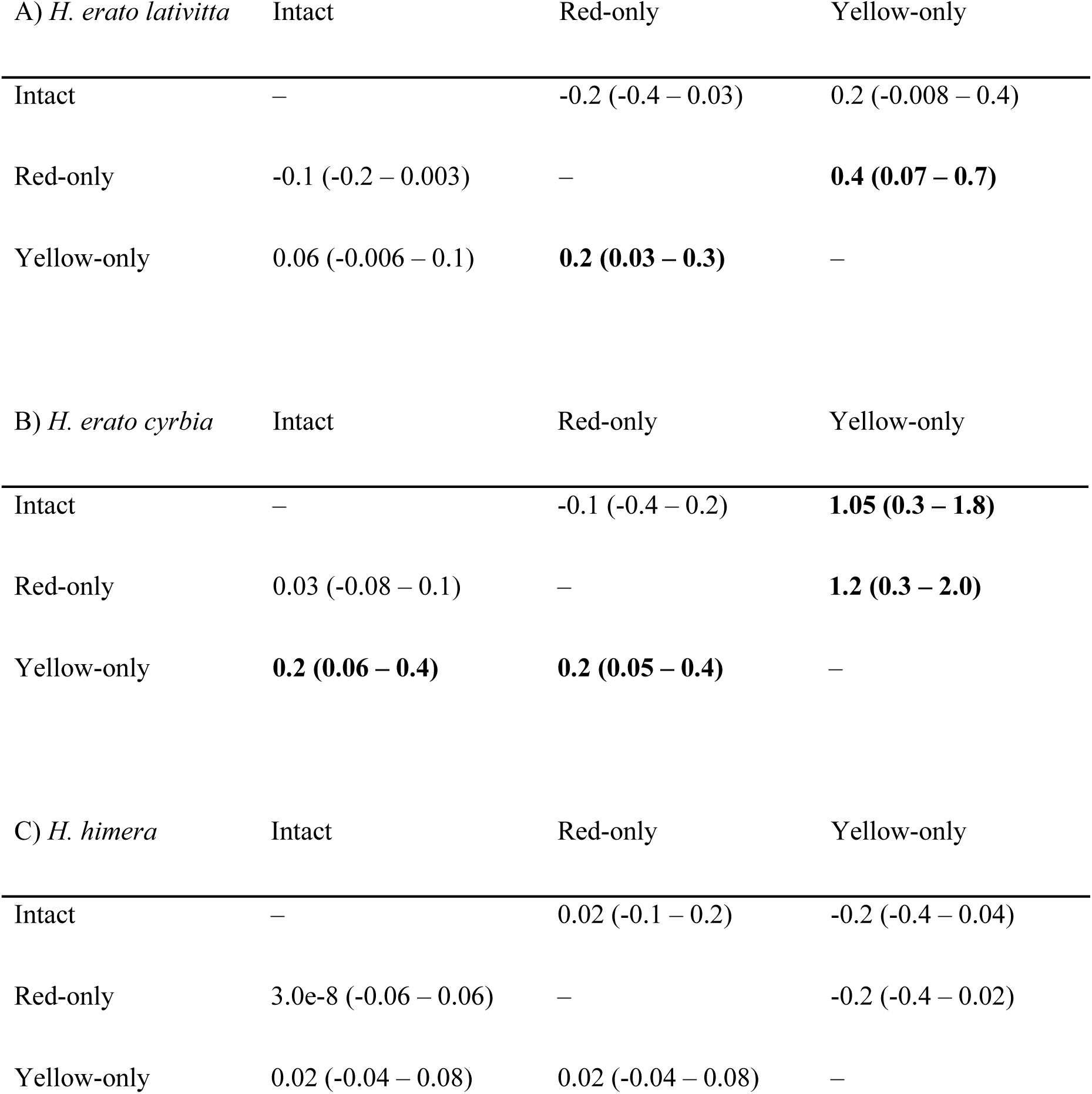
Contrast estimates showing effect size and direction of differences in estimated marginal means of preference behaviors of A) *H. erato lativitta* males (n = 21), B) *H. erato cyrbia* males (n = 15) and C) *H. himera* males (n = 16). Rows separate between preference for the three different female wing pattern models: the ‘Intact’ model was the full *H. erato lativitta* wing pattern; ‘Red-only’ was the model with only the red pattern elements of the *H. erato lativitta* wing pattern; ‘Yellow-only’ was the model with only the yellow pattern elements of the *H. erato lativitta* wing pattern (see Figure 2A-C). Values indicate difference in probability of approach (above diagonal, rows minus columns) or courtship (below diagonal, columns minus rows) with 95% confidence intervals. Values in bold indicate significant differences (p < 0.05) for each population in their ‘approach’ or ‘courtship’ towards the compared model types.

## Data Availability

Analyses reported in this article can be reproduced using the data provided by Smith et al. (2024).

## Results

In total, we measured preferences of 21 *H. erato lativitta*, 15 *H. erato cyrbia* and 16 *H. himera* males over 122 behavioral trials. The three male types responded differently to the female models, both in terms of the number of ‘approaches’ and number of ‘courtships’ (Figure 3). Model testing revealed a significant reduction in deviance when the interaction between male species and model type was removed from our GLMMs for both ‘approach’ (LRT: 2ΔlnL = 140.45, d.f. = 4, p < 2.2e^-16^) and ‘courtship’ (LRT: 2ΔlnL = 43.3, d.f. = 4, p = 9.06e^-9^) indicating that the males respond differently to different pattern elements. Weather strongly predicted male ‘approach’ behaviors (LRT: 2ΔlnL = 11.8, d.f. = 3, p < 0.01) but not ‘courtship’ behaviors (LRT: 2ΔlnL = 7.6, d.f. = 3, p = 0.06), however, there was no significant difference between the EMM of male approach probability towards each model in each weather category (SM Figures 1-3; SM Table 2). Nevertheless, we retained weather in all models because it influenced the likelihood of a male approaching the model, and all courtship behaviors were preceded by a male approaching a model. For both ‘approach’ and ‘courtship’ the minimum adequate model included the male species*model type interaction term, as well as male species, model type and weather as fixed effects. To further investigate differences in preference for wing pattern across the three species, we calculated the EMM of proportion of minutes with ‘approaches’ and ‘courtships’ towards each model type separately for each species. *H. erato lativitta* did not distinguish between the intact and red-only model but was more likely to ‘approach’ and ‘court’ the intact and red-only model than the yellow-only model (Table 1A). Looking across all three male types, we found similar preference (both in terms of ‘approach’ and ‘courtship’) between *H. erato lativitta* and *H. erato cyrbia*; both preferred the intact and red-only models most, and the yellow-only model least, while *H. himera* showed no model preference in either ‘approach’ or ‘courtship’ behaviors (Table 1).

**Figure 3.**
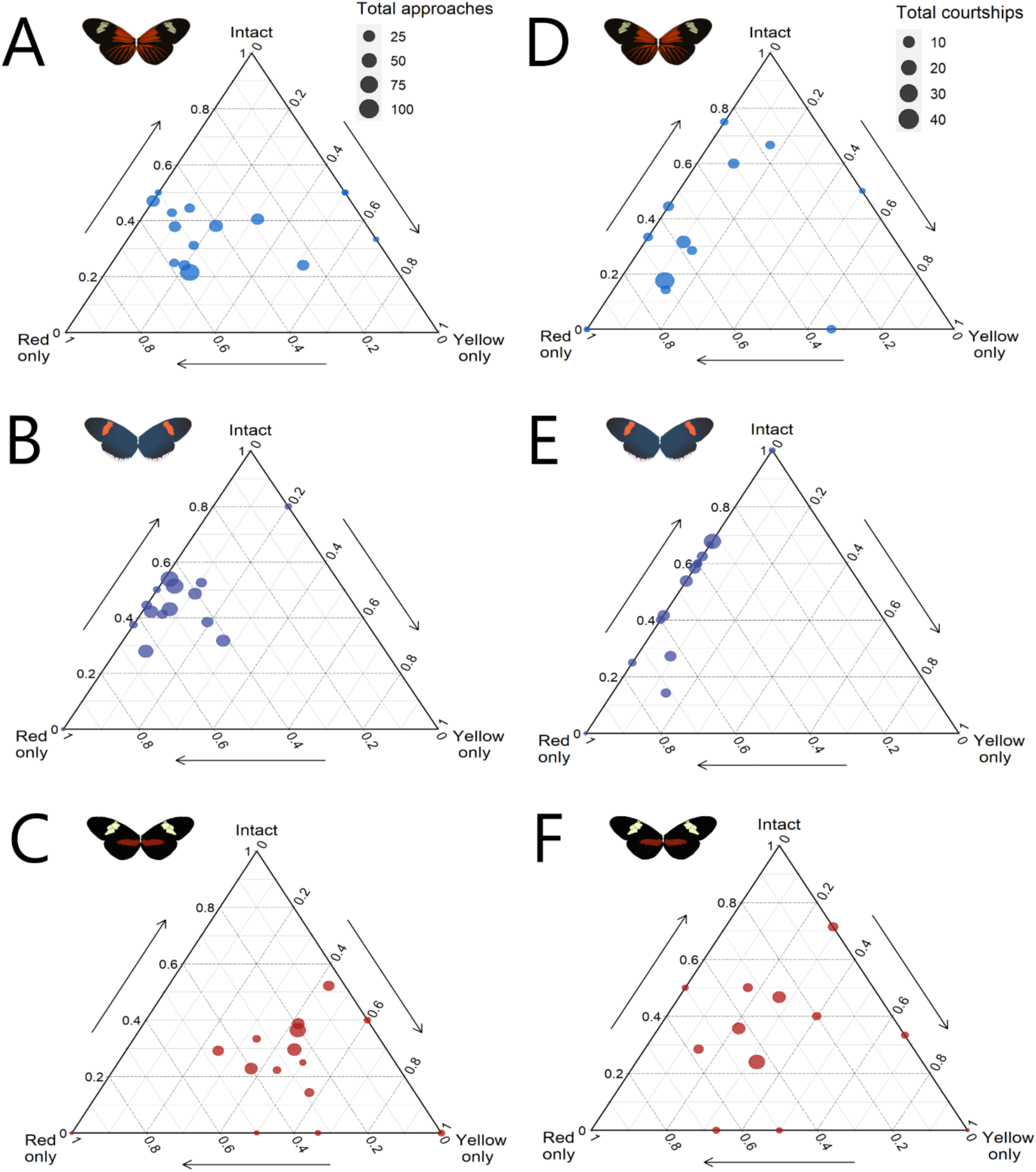
Ternary plots showing the proportion of minutes in which individual males directed preference behaviors towards either of the three female wing pattern models. A-C shows approach behaviors in (A) *H. erato lativitta* (n = 21, mean total approaches per male = 19.3), (B) *H. erato cyrbia* (n = 15, mean total approaches per male = 26.6), and (C) *H. himera* (n = 16, mean total approaches per male = 20.4). D-F shows hovering courtship behaviors in (D) *H. erato lativitta* (n = 21, mean total courtships per male = 3.9), (E) *H. erato cyrbia* (n = 15, mean total courtships per male = 5.1), and (F) *H. himera* (n = 16, mean total courtships per male = 3.8). Left ternary axis shows the proportion of minutes in which males directed preference towards the intact model, bottom ternary axis, towards the red only model, and right axis towards the yellow only model. Each point represents the mean responses of males across all trials, and point size represents the cumulative activity level of that male across all trials. Males that did not show any courtship responses are not represented in the ternary plots. Butterfly images in the top left of each plot show the wing pattern of the male species of the plot.

## Discussion

We found that experimentally removing individual warning color pattern elements significantly influenced male behaviors; however, this effect differed between males of closely related species. For *H. erato lativitta*, individual color pattern elements influenced male courtship response, but the full pattern model was not preferred most of all. Male *H. erato lativitta* preferred to court the model with the red dennis-ray pattern only over the model that lacked any red elements, but there was no difference in preference between the intact wing pattern model and the model without the yellow forewing element. This suggests that the red dennis-ray elements are more attractive to male *H. erato lativitta* than yellow elements, and that this part of the pattern may act independently in signaling to courting males. At first, these results might seem surprising: the survival of these butterflies relies heavily on predators recognizing their wing patterns as toxic, reinforced by mimicry between co-occurring species. *H. erato lativitta* is part of a mimicry ring with several *Heliconius* species found in the Ecuadorian Amazon, as well as some non-*Heliconius* and even some moths, all of which have both the red dennis-ray pattern and yellow patch in their warning pattern (Jiggins 2016). This would predict strong selection for males to prefer females with this shared warning pattern, to maintain the recognized signal in their offspring.

The lack of strong preference in *H. erato lativitta* for females with all the components of its own color pattern shows that some elements may be redundant to the *H. erato lativitta* visual mating signal. This may indicate that there is variation in the exact information communicated across the wing. These findings contribute to the collection of studies showing that individual wing pattern elements serve a variety of functions, and not all are used in mate preference (Candolin 2003). For example, in a study of plumage evolution in hummingbirds, Beltrán and colleagues proposed that females receive different information from different plumage patches; those which are similar across species could be used for mate quality evaluation, while patches which differ were more likely to be used in species discrimination (Beltrán et al. 2021). We cannot speak to the exact functions of individual *H. erato lativitta* pattern elements, but the results presented here indicate that individual wing pattern elements can act independently as signals.

When comparing among species of *Heliconius*, it appears that most of the differences were associated with presence or absence of red pattern elements. Despite very different wing patterns, both *H. erato* subspecies were similar in approaching the yellow-only model significantly less than the intact or red-only models, while *H. himera* showed no model preference. This difference is noteworthy considering evidence of premating isolation between *H. himera* and *H. erato cyrbia* based on wing pattern (Merrill et al. 2014). *H. himera* and *H. erato cyrbia* are sister species, in which assortative mating is maintained at a rate of 90% across their hybrid zone at least in part by visual premating isolation (Jiggins et al. 1996, Merrill et al. 2014). The *H. himera* wing pattern features a yellow forewing patch, similar in appearance to the yellow patch on the *H. erato lativitta* pattern, while *H. erato cyrbia* features a red forewing patch and no yellow forewing elements (Figure 2). It could be that *H. erato cyrbia* males preferentially avoid the yellow-only model because it gave the same mating signal as the wing pattern of *H. himera*. However, *H. himera* males did not show preference for the models with the yellow forewing patch, despite this similarity. In fact, *H. himera* males exhibited no differences in preference between any of the models, which could be driven by lower investment in visual sensitivity in *H. himera* compared to *H. erato* (Montgomery & Merrill 2017). Dell’Aglio et al. (2022) similarly found that while *H. himera* does discriminate based on visual cues, olfactory cues were weighted more strongly in a foraging assay. In the context of mate choice, the lack of *H. himera* preference may also be due to the local butterfly community: *H. erato cyrbia* co-occurs with other *Heliconius* species, including co-mimics with similar blue iridescent patterns, whereas the *H. himera* habitat contains comparatively fewer *Heliconius*, with no direct co-mimics (Jiggins et al. 1996). Thus, there may be stronger selective pressure for *H. erato cyrbia* to visually discriminate between species. Increased preference for red elements in *H. erato* may also be driven by sensory bias; red is a common color of many flower species that *Heliconius* feed on, and they are known to have an innate preference for red/orange (Merrill et al. 2011).

However, this preference has been shown to be weaker in *H. himera* than *H. erato cyrbia*, which may be due to more diverse flower colors in the *H. himera* habitat (Dell’Aglio et al. 2022). It is striking to find that male *H. erato cyrbia* showed preference for red wing pattern elements even in a very different wing pattern, however this phenomenon has also been observed in Lycaenid butterflies (Fordyce et al. 2002). Although subtle, these differences may demonstrate that premating isolation between *H. erato cyrbia* and *H. himera* is maintained by preference in *H. erato cyrbia* for red forewing elements, and lack of visual preference in *H. himera*.

We also found no overall significant differences in preference between weather conditions, although weather did interact with model type in our statistical model of ‘approach’ behavior. This is in contrast to Hausmann et al. (2021), who found that *Heliconius* butterflies are more likely to court their own color pattern with brighter light. Across our study, over 80% of our trials occurred in overcast or highly cloudy conditions, therefore it is likely that we did not experience enough days of bright light while conducting our trials to detect strong differences in preference. This could also explain why we did not detect strong preference differences, as the low light conditions resulted in less active males, fewer courtships, and less strong discrimination between wing patterns models by males.

Visual mate choice signals play a role in the early stages of divergence in *Heliconius* butterflies (Merrill et al. 2015). It would seem that *H. himera* has diverged from the two *H. erato* included in this study, displaying weaker visual preferences based on wing pattern components. *H. erato cyrbia,* meanwhile, possesses a very different wing pattern to *H. erato lativitta*, yet male preference for specific wing pattern elements is maintained. This hints as to the process of speciation with wing pattern divergence in these species; while wing pattern divergence across all *Heliconius* taxa is driven by selection for mimicry (Bates 1862), male preference can concurrently diverge to exploit key preference elements that are maintained, rather than a complete overhaul of preference to a fully new wing pattern. For species such as *Heliconius* butterflies, whose morphological evolution is constrained by selection for aposematic warning signals, this illuminates how mate choice based on wing pattern can persist as wing patterns diversify. By enacting mating preferences on only particular elements of the mating signal, selection for aposematic warning patterns and the function of wing patterns as mating cues are balanced across the full wing pattern. The differences in preference between *H. erato* and *H. himera* illustrate this idea, showing that strong differences in preference for colors of the forewing pattern may be enough to drive assortative mating. Previous studies have found that mate preferences in *Heliconius* are commonly restricted to discrimination between forewing patterns (Jiggins et al. 2001, Kronforst et al. 2006). This releases the hind wing pattern from the constraining selection of mate preference and allows the pattern to be more flexible to selection by aposematism and mimicry. Whether predators impose differential selection pressures on different wing elements, or different wings, is yet to be explored in *Heliconius*.

## Supplementary Materials

We tested for the relationship between our four weather variables (‘100% overcast’, ‘>50% overcast’, ‘<50% overcast’ and ‘clear’) and temperature, and the effect of these variables on both approach and courtship. First, to test how temperature and weather relate, we fitted a linear regression of temperature against the categorical variables of weather, and then tested the significance of this relationship in an ANOVA. Overall, 51% of trials occurred in overcast weather, 32% occurred in high cloud weather, 11% in low cloud weather and 6% in clear weather. The mean temperature of the trials was 27.5°C (SD = 4.1°C), and average temperature decreased with increasing cloud cover (SM Figure 1; one-way ANOVA: F = 493.15, D.F. = 3, p < 2.2e^-16^). This showed that temperature and weather are highly correlated, so then we tested how male activity level varied with weather category. For this, we used the binomial counts of approach or courtship behavior as in our analysis of preference differences, but fitted these separately in a GLMM with only weather as an independent variable. We then used LRT to test for a significant difference in deviance when weather was removed, and estimated the EMMs of the activity level across each weather category.

**SM Table 1.** Results of model optimization comparisons.

| Dependent variable | Independent variables | Random effects | Log likelihood of model optimizers |  |  |  |  |
| --- | --- | --- | --- | --- | --- | --- | --- |
|  |  |  | Nelder-Mead | bobyqa | nlminbwrap | nloptwrap.<br>NLOPT_L<br>N_NELDE<br>RMEAD | nloptwrap.<br>NLOPT_L<br>N_BOBY<br>QA |
| Approach | weather +<br>model_type *<br>male_species | (1 \| male_id) + (1 \|<br>model_set/trial_id) | -1579.724 | -1579.724 | -1579.724 | -1579.724 | -1579.724 |
| Approach | weather +<br>model_type +<br>male_species | (1 \| male_id) + (1 \|<br>model_set/trial_id) | -1649.948 | -1649.948 | -1649.948 | -1649.948 | -1649.948 |
| Approach | weather +<br>model_type | (1 \| male_id) + (1 \|<br>model_set/trial_id) | NA | NA | NA | NA | NA |
| Approach | weather +<br>male_species | (1 \| male_id) + (1 \|<br>model_set/trial_id) | NA | NA | NA | NA | NA |
| Courtship | weather +<br>model_type *<br>male_species | (1 \| male_id) + (1 \|<br>model_set/trial_id) | -651.8558 | -651.8541 | -651.8541 | -651.8541 | -651.8541 |
| Courtship | weather +<br>model_type +<br>male_species | (1 \| male_id) + (1 \|<br>model_set/trial_id) | -673.4944<br>8 | -673.4937 | -673.4937 | -673.4937 | -673.4937 |
| Courtship | weather +<br>model_type | (1 \| male_id) + (1 \|<br>model_set/trial_id) | -674.4726 | -674.4705 | -674.4705 | -674.4705 | -674.4705 |
| Courtship | weather +<br>male_species | (1 \| male_id) + (1 \|<br>model_set/trial_id) | NA | NA | NA | NA | NA |

**SM Table 2.** Results of multiple comparison test of effect of weather categories on ‘approach’ behaviors.

| Contrasting weather categories | Estimate of difference | SE | Lower confidence interval | Upper confidence interval | P-value |
| --- | --- | --- | --- | --- | --- |
| Clear – over 50 % cloud | 0.1 | 0.3 | -0.7 | 0.9 | 1.0 |
| Clear - overcast | 0.4 | 0.3 | -0.4 | 1.2 | 0.5 |
| Clear – under 50% cloud | 0.03 | 0.4 | -0.9 | 1.0 | 1.0 |
| Over 50% cloud - overcast | 0.3 | 0.14 | -0.03 | 0.7 | 0.08 |
| Over 50% cloud – under 50% cloud | -0.07 | 0.24 | -0.7 | 0.57 | 1.0 |
| Overcast – under 50% cloud | -0.4 | 0.24 | -1.0 | 0.2 | 0.3 |

**SM Figure 1.**
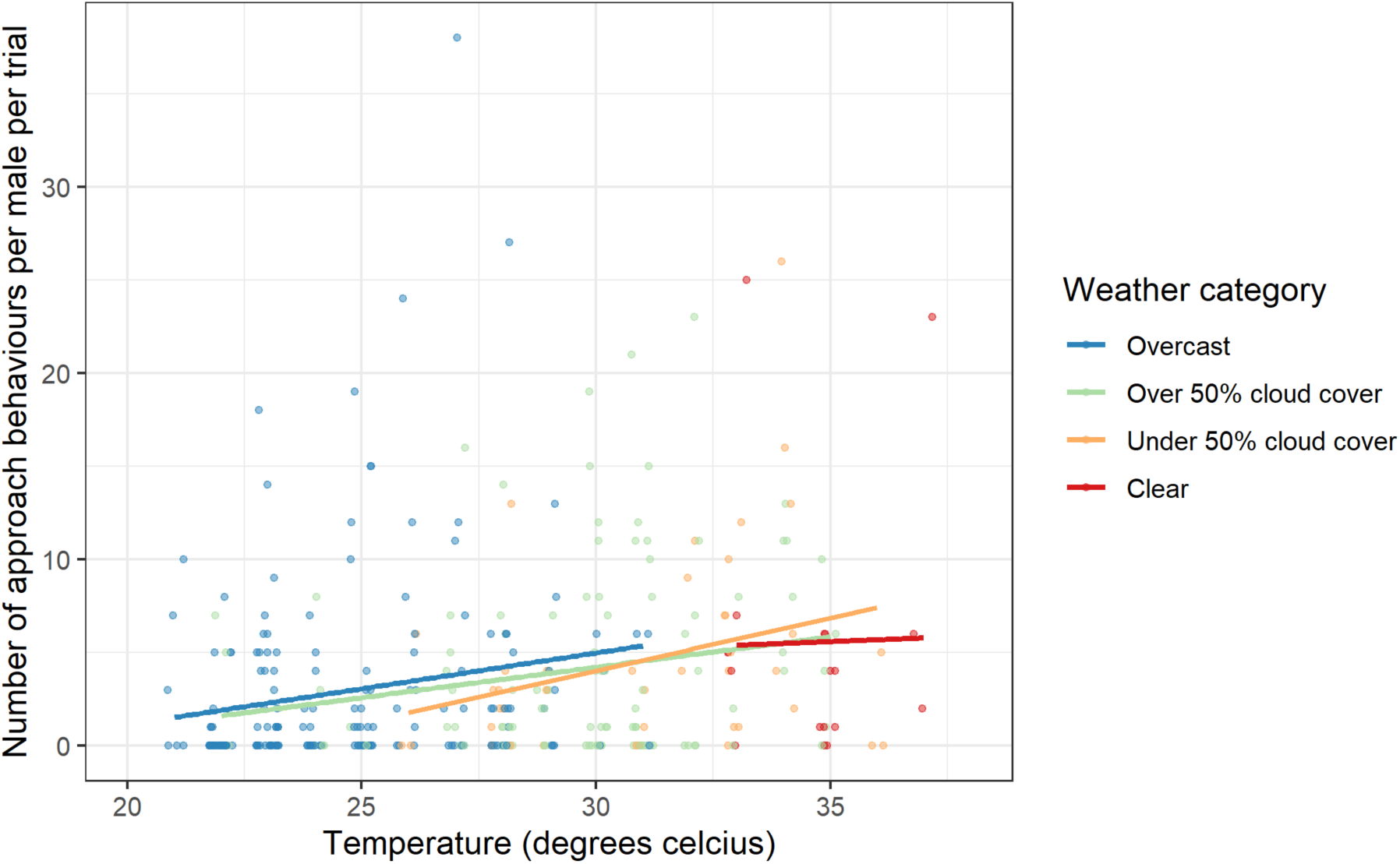
Number of approach behaviors recorded per male per trial, per weather category, plotted against the temperature recorded during the trial. Points are colored by the weather category the trial occurred in, and a line of best fit for each weather category is fitted to the graph.

**SM Figure 2.**
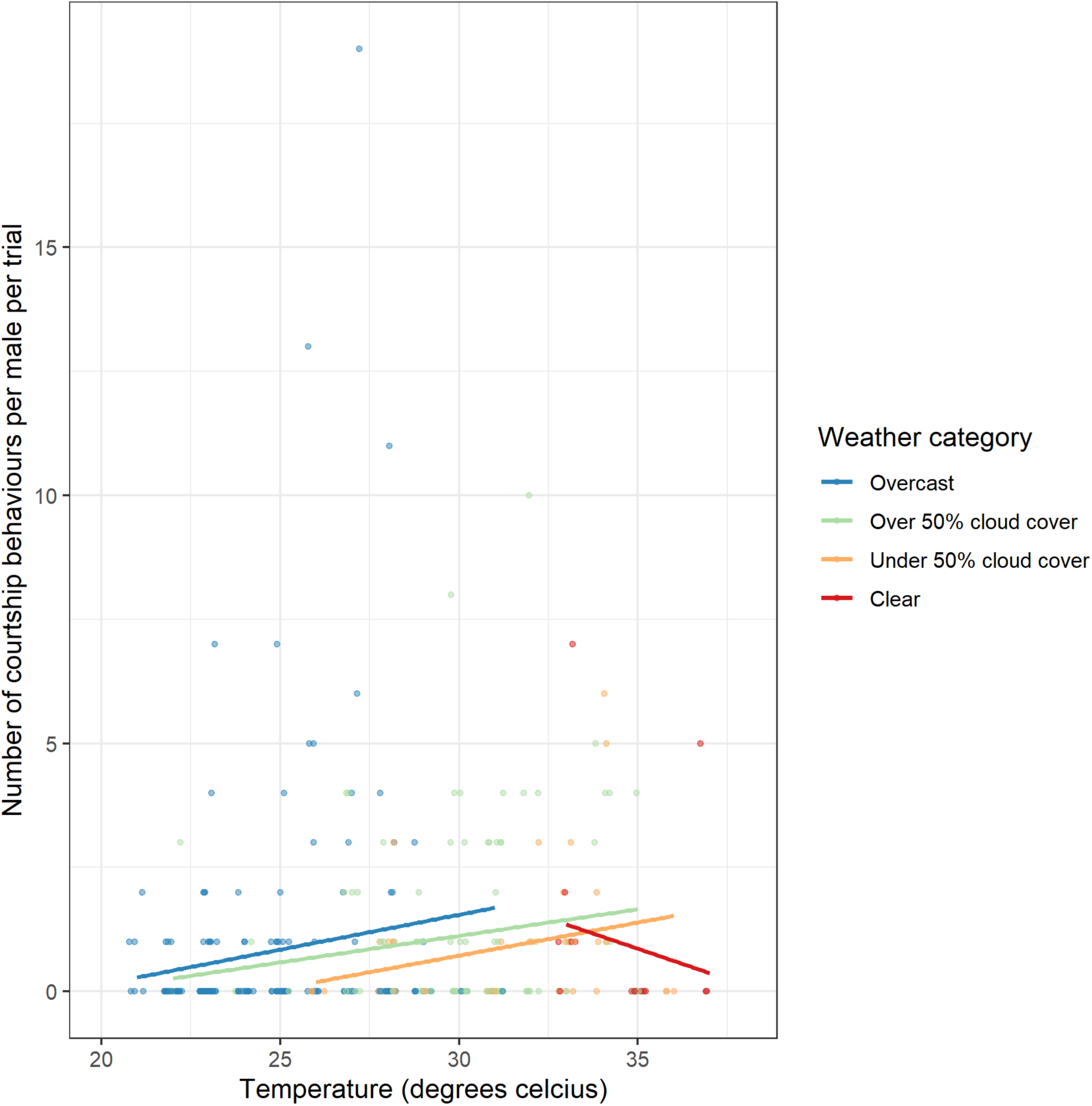
Number of courtship behaviors recorded per male per trial, per weather category, plotted against the temperature recorded during the trial. Points are colored by the weather category the trial occurred in, and a line of best fit for each weather category is fitted to the graph.

**SM Figure 3.**
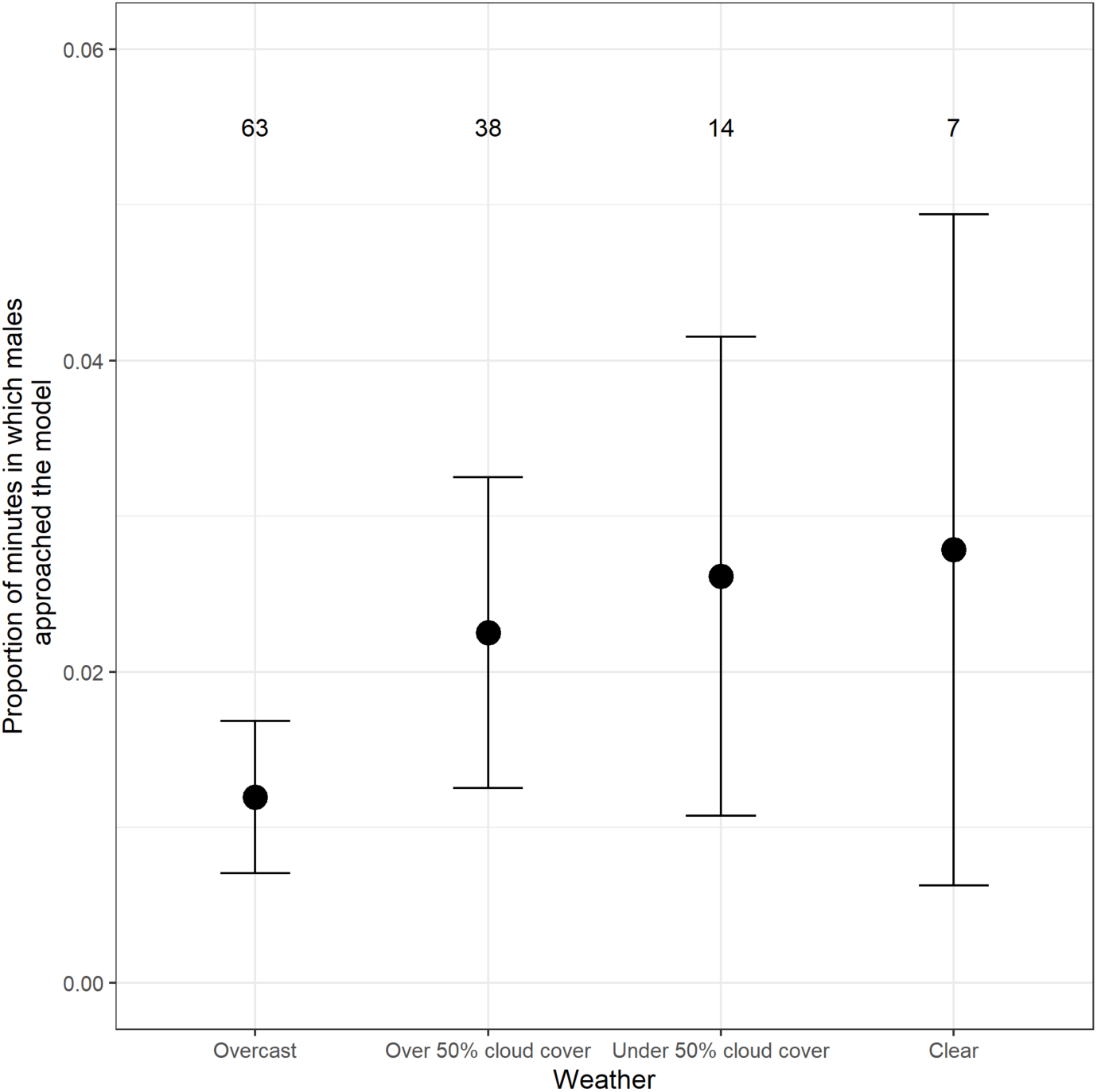
Approach behavior of males during trials of each weather category, shown by the estimated marginal means of proportion of minutes during the trial in which males of all species approached the models. Error bars show the 95% confidence intervals, based on the analysis. Numbers above the error bars show the number of trials conducted in each weather category.

